# LILRB3 (ILT5) is a myeloid checkpoint on myeloid cells that elicits profound immununomodulation

**DOI:** 10.1101/2020.06.04.135400

**Authors:** Muchaala Yeboah, Charys Papagregoriou, Des C. Jones, H.T. Claude Chan, Guangan Hu, Justine S. McPartlan, Torbjörn Schiött, Ulrika Mattson, C. Ian Mockridge, Ulla-Carin Tornberg, Björn Hambe, Anne Ljungars, Mikael Mattsson, Ivo Tews, Martin J. Glennie, Stephen M. Thirdborough, John Trowsdale, Björn Frendeus, Jianzhu Chen, Mark S. Cragg, Ali Roghanian

## Abstract

Despite advances in identifying the key immunoregulatory roles of many of the human leukocyte immunoglobulin (Ig)-like receptor (LILR) family members, the function of the inhibitory molecule LILRB3 (ILT5, CD85a, LIR3) remains unclear. Studies indicate a predominant myeloid expression; however, high homology within the LILR family and a relative paucity of reagents have hindered progress for this receptor. To investigate its function and potential immunomodulatory capacity, a panel of LILRB3-specific monoclonal antibodies (mAb) was generated. LILBR3-specific mAb bound to discrete epitopes in either Ig-like domain two or four. LILRB3 ligation on primary human monocytes by agonistic mAb resulted in phenotypic and functional changes, leading to potent inhibition of immune responses *in vitro*, including significant reduction in T cell proliferation. Importantly, agonizing LILRB3 in humanized mice induced tolerance and permitted efficient engraftment of allogeneic cells. Our findings reveal powerful immunosuppresive functions of LILRB3 and identify it as an important myeloid checkpoint receptor.

## Introduction

Molecules of the human leukocyte immunoglobulin (Ig)-like receptor (LILR) family, discovered over two decades ago (*1, 2*), are expressed on leukocytes and are commonly dysregulated in a wide range of pathologies (*3–5*). There are five activating (LILRA1, 2, 4–6), five inhibitory (LILRB1–5) and one soluble (LILRA3) LILR that together regulate immune responses (*3*). They display two, or four, homologous C-2 type Ig-like extracellular domains, but differ in their transmembrane and cytoplasmic regions (*2, 6*). LILRA have short truncated cytoplasmic tails with charged arginine residues in their transmembrane domains, facilitating association with the ITAM-bearing FcεR γ-chain to propagate activating signaling cascades (*7*). Conversely, LILRB have long cytoplasmic tails that contain multiple ITIM motifs, which recruit phosphatases such as SHP-1 and SHIP-1 to elicit inhibitory signaling (*2, 6*). Located at human chromosome 19q13.4, these receptors demonstrate significant allelic variation, with LILRB3, LILRB4 (ILT3) and LILRA6 (ILT8) each displaying at least 15 different variants (*2, 8–10*).

The LILRB molecules are proposed to act as immune checkpoints serving to control and limit overt immune responses (*3*). In agreement with this, LILRB expression is increased in suppressive (also referred to as alternatively activated or M2) macrophages and tolerogenic dendritic cells (DCs) (*11–15*). On monocytes, co-ligation of LILRB1 (ILT2) and LILRB2 (ILT4) with the activatory FcγRI (CD64) results in SHP-1 activation, decreasing downstream phosphorylation events and intracellular calcium mobilization (*16*). Engagement of LILRB1 on macrophages by the common HLA-I subunit, β_2_-microglobulin, on malignant cells limits their phagocytic potential (*17*). Similarly, we and others have shown that ligation of LILRB1, 2 or 4 renders DCs tolerogenic, leading to inhibition of T cell responses (*11, 12, 15, 18–21*). As such, the engagement of LILRB1 and LILRB2 by their high affinity ligand HLA-G is an important immunosuppressive pathway at the fetal-maternal interface during pregnancy (*22–24*) and may also be involved in tumor immunoevasion (*5*).

Although mice do not express LILRs, they possess an orthologous system comprised of two paired Ig-like receptors (PIR); the activating PIR-A and the inhibitory PIR-B. PIR-B regulates priming of cytotoxic T-lymphocytes by DCs through interaction with MHC class I (MHCI) (*25*) and negatively influences integrin signaling in neutrophils and macrophages (*26*). Furthermore, PIR-B regulates the differentiation of myeloid-derived suppressor cells (MDSCs) that aid tumor progression (*27*).

Among the inhibitory LILRBs, LILRB3 (ILT5/LIR3/CD85a), containing 4 extracellular Ig-like domains and 4 intracellular ITIM motifs, represents an attractive immunomodulatory target because of its relative restriction to, and high expression on, myeloid cells (*3, 4*). However, due to the lack of specific reagents and model systems, its exact functions and immunoregulatory potential have not been fully explored. In this study, we addressed this by generating a bespoke panel of novel LILRB3-specific mAb, some of which were used to probe the function of LILRB3 in relevant preclinical platforms. Our data demonstrate that LILRB3 activation confers potent immunoinhibitory functions through reprograming and tolerizing of myeloid cells, and suggest that modulating LILRB3 activity may provide exciting new treatment strategies in various disease settings, such as transplantation.

## Results

### Generation and characterization of a panel of fully human LILRB3-specific mAb

To study the protein expression and function of LILRB3, LILRB3-specific antibodies were identified from a human antibody phage-display library, n-CoDeR (*28, 29*). Initial alignment analysis of extracellular domains of LILRB1–5 indicated the presence of a limited number of conserved amino acid (a.a.) residues across the LILRB3 ectodomain (fig. S1), against which specific mAb could be generated. In this regard, phages binding to the ‘target’, ectodomain of LILRB3 protein (present in solution, coated to a plastic surface or expressed on cells), and not to the homologous (~65% extracellular homology) ‘non-target’ LILRB1 ectodomain protein, were selected (Fig. 1A; fig. S2). To increase specificity and yield, the cross-reactive phages were initially removed through a pre-selection (negative selection/depletion using ‘non-target’ proteins), followed by the selection itself (positive selection). Following each selection round, the selected clones were screened against the ectodomains of both LILRB1 and LILRB2 by flourometric microvolume assay technology (FMAT) and enzyme-linked immunosorbent assay (ELISA), and cross-reactive clones were further excluded from the panel. After three rounds of phage panning and enrichment, successful selection of clones specific for LILRB3 was reconfirmed by FMAT and ELISA, with target-specific phage converted to soluble scFv and screened further (Fig. 1B and C). Successful clones were selected based on binding to LILRB3 and lack of cross-reactivity to LILRB1 and LILRB2. Selected scFv clones (>200) were then sequenced and tested for binding against primary cells and LILRB transfectants using high throughput flow cytometry (Fig. 1D). Subsequently, 46 candidate target-specific clones were converted to human IgG1 (hIgG1) and, in addition to screening against LILRB1–3 transfectants, to exclude those with potential broader LILR cross-reactivity, were screened against a larger panel of LILR-expressing cell lines (Fig. 1E). Due to cross-reactivity to one or more other LILR family members, as exemplified by clone A30 (Fig. 1E, bottom panel), 30 mAb clones were further excluded at this stage. In total a panel of 16 LILRB3-specific antibodies were identified for further study. These LILRB3-specific clones were further tested and confirmed to have no cross reactivity to the mouse orthologue, PIR-B (data not shown). A selection of these mAb were then fluorochrome-labelled and used to determine the LILRB3 expression profile of human peripheral blood; demonstrating predominant staining of monocytes and to a lesser extent granulocytes (Fig. 1F and G), in agreement with previous reports (*2, 3, 6*). The immunophenotyping also revealed that LILRB3 expression was significantly higher on circulatory CD14^+hi^/CD16^−^ ‘classical’ and CD14^+^/CD16^+low^ ‘intermediate’ monocytes compared with the more inflammatory CD14^+^/CD16^+hi^ ‘non-classical’ monocytes (Fig. 1F and G).

**Fig. 1.**
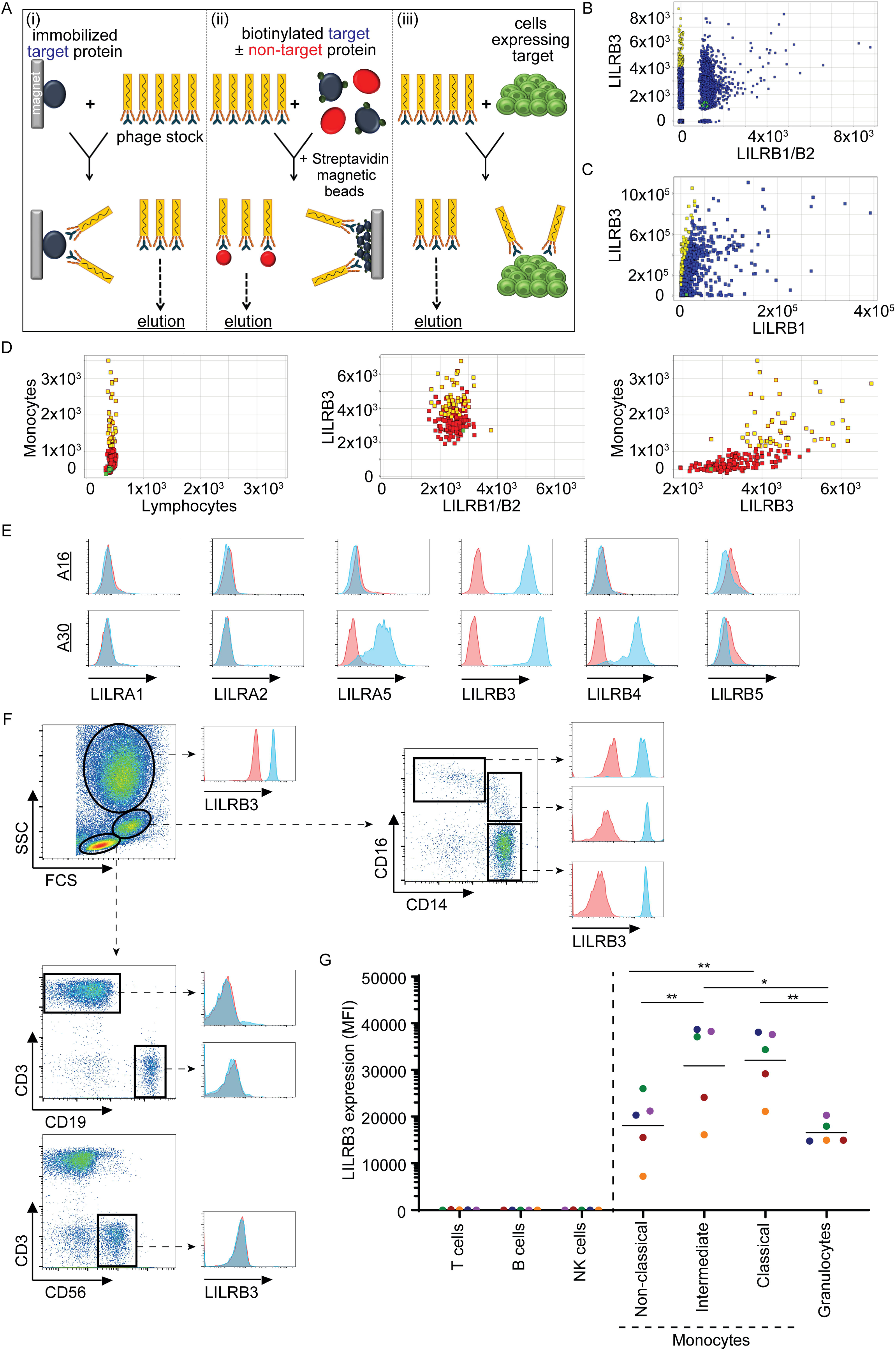
Generation of fully human mAb against LILRB3. (**A**) Schematic of antibody generation by phage display via three independent ‘panning’ techniques; *(i)* immobilized target (LILRB3), *(ii)* biotinylated target and excess non-target (LILRB1), and *(iii)* LILRB3-transfectant cell lines (from left to right). Biopanning was performed against generated target protein using a scFv library; ‘non-target’ cross-reactive scFv clones were eluted off at each panning stage and target-specific scFv clones were then converted to a soluble format, sequenced and screened by various cell- and protein-based assays. (**B-C**) Screening of generated LILRB3 clones. (**B**) FMAT and (**C**) ELISA were performed and scFv clones screened against LILRB3 target and LILRB1/LILRB2 non-target transfected CHO-S cells and extracellular LILRB1 protein, respectively. MFI of binding to each target was calculated, with target-specific scFv depicted in yellow, non-target scFv in blue and the irrelevant isotype control shown in green. (**D**) Screening of LILRB3 scFv clones by high throughput flow cytometry. PBMCs or LILR-transfected CHO-S cells were incubated with His-tagged scFv supernatants, followed by secondary anti-His staining. Where transfected CHO-S cells were used, LILRB1- and LILRB2-transfected cells were used as non-targets for LILRB3. Clones were compared against both gated CD14^+^ monocytes and target transfected CHO-S cells, with LILRB3 specific clones highlighted in yellow, non-specific clones in red and the isotype control in green. (**E**) Specificity of LILRB3 clones against human LILR-transfected 2B4 cells. LILRB3 mAb were tested against cells stably transfected with the indicated LILR family members by flow cytometry; a representative specific clone (A16; top panel) and a non-specific cross-reactive clone (A30; bottom panel) shown. (**F-G**) Testing the specificity of directly fluorochrome-labeled LILRB3 clones against primary cells by flow cytometry. (**F**) Fresh whole peripheral blood stained with either APC-labelled LILRB3 (represented by clone A16) or an irrelevant human (h) IgG1 isotype control as well as various leukocyte surface markers, as indicated. Dot plots and histograms are representative of multiple donors indicating gating of each leukocyte subset as indicated: T cells, B cells and NK cells, monocytes and granulocytes. (**G**) Graph showing relative expression of LILRB3 on each leukocyte subset. Two-tailed paired T-test performed (* *p* < 0.05; ** *p* < 0.005); *n* = 5 independent donors (each color represents an individual donor).

The selected LILRB3 mAb were also tested for their specific binding properties. Surface plasmon resonance (SPR) analysis showed that all LILRB3-specific clones bound to recombinant LILRB3-hFc protein in a dose-dependent manner (as represented by A16; Fig. 2A) and displayed a range of affinities (Table 1). Interestingly, all mAb had similar association rates (~10^5^), but varied in their dissociation rates by three orders of magnitude (~10^−3^−10^−6^).

**Table 1.** LILRB3 mAb characterization. Summary of binding domain, blocking potential and affinity measurements for the selected LILRB3-specific mAb.

| Clone ID | Ig-like domain | Blocking potential | Affinity ($K_D$) |
| --- | --- | --- | --- |
| **1 (A1)** | 4 | N | $3.12 \times 10^{-10}$ |
| **2 (A12)** | 2 | Y | $5.30 \times 10^{-10}$ |
| **3** | 4 | N | $2.92 \times 10^{-10}$ |
| **4 (A16)** | 4 | N | $2.88 \times 10^{-9}$ |
| **5** | 2 | Y | $1.04 \times 10^{-9}$ |
| **6** | 2 | N | $1.55 \times 10^{-9}$ |
| **7** | 4 | N | $3.85 \times 10^{-10}$ |
| **8 (A28)** | 4 | N | $1.46 \times 10^{-9}$ |
| **9** | 4 | N | $9.84 \times 10^{-9}$ |
| **10** | 2 | N | $3.71 \times 10^{-10}$ |
| **11** | 4 | N | $1.37 \times 10^{-9}$ |
| **12** | 4 | N | $1.24 \times 10^{-10}$ |
| **13** | 4 | N | $3.25 \times 10^{-9}$ |
| **14** | 2 | N | $3.05 \times 10^{-10}$ |
| **15** | 4 | N | $1.56 \times 10^{-9}$ |
| **16** | 2 | N | $1.68 \times 10^{-8}$ |
| **222821** | 2 | - | $1.47 \times 10^{-11}$ |

Clone 222821 represents the commercial LILRB3 mAb (mIgG2a).
Ability to block binding of clone 222821 is indicated by Yes (Y) or No (N).
Affinity assessed using the univalent model of 1:1 binding by SPR; clone IDs described herein are indicted in brackets.

**Fig. 2.**
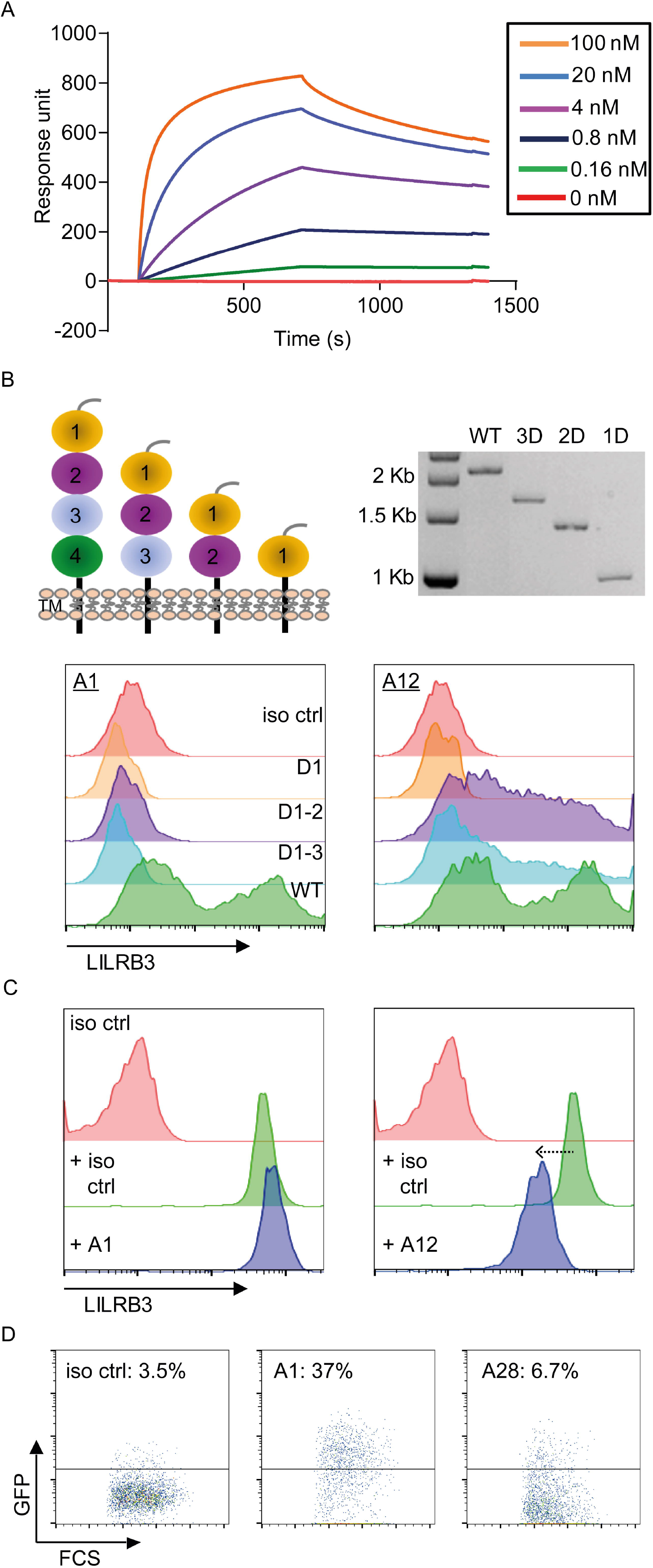
Characterization of LILRB3 antibodies. (**A**) LILRB3 mAb affinity assessed by SPR. LILRB3-hFc recombinant protein was immobilized and various LILRB3 mAb flowed across the chip. Representative LILRB3 clone A16 shown. (**B**) LILRB3 domain epitope mapping. HEK293F cells transfected with either WT LILRB3 (full-length extracellular portion), D1-3, D1-2 or D1 were stained with LILRB3 clones, followed by an anti-hIgG secondary antibody. Schematic of domain constructs and restriction digest of each DNA construct shown (top panel). Histograms showing staining of two representative clones differentially binding to color-coded WT (D4), D1-3, D1-2 and D1-expressing cells (bottom panel; *n* = 3 independent experiments). (**C**) Ability of generated mAb to cross-block binding of a commercial LILRB3 mAb (clone 222821). PBMCs were stained with unconjugated LILRB3 antibody clones and subsequently stained with a directly-conjugated 222821 mAb and analyzed by flow cytometry; representative clones displayed (A1 non-blocking; A12 partial blocking), as indicated. (**D**) LILRB3 2B4 reporter cells were incubated with 10 μg/ml LILRB3 antibodies overnight to assess receptor signaling potential as judged by GFP induction measured by flow cytometry; representative clones with percentage of GFP expression shown.

Cell surface epitope mapping studies were then performed and compared to a commercial mAb (clone 222821), using a series of LILRB3 extracellular domain (D) mutants displaying either all four Ig domains (wild-type [WT]), three, two, or one domain, transiently transfected into HEK293F cells. Two distinct groups of mAb were identified: those that bound to the WT, D3 and D2 expressing cells (including clone 222821 and exemplified by A12); and those that bound only the WT-transfected cells (exemplified by A1) (Fig. 2B). Although conserved a.a. residues are present throughout the ectodomain (fig. S1), the selected mAb were shown to bind either within D2 or D4 (6/16 and 10/16 clones, respectively; Table 1), perhaps indicating improved accessibility for these regions within the 3D structure. In agreement with this, subsequent blocking assays confirmed that a number of D2-binding mAb reduced the binding of the commercial mAb (*e.g.*, A12), suggesting a shared or related epitopes; whilst others did not (*e.g.*, A1), confirming binding to discrete epitopes (Fig. 2C and Table 1).

Subsequently, reporter cells transfected with a chimeric receptor expressing the extracellular domain of LILRB3, fused with the human CD3ζ cytoplasmic domain, were used to investigate whether the generated mAb were able to crosslink the receptor. Cross-linking results in the production of NFAT activation and the subsequent expression of GFP and is indicative of agonistic potential (*30*). Using these cells, we were able to identify two distinct groups of LILRB3 mAb, those with ‘agonistic’ activity capable of inducing signaling upon binding to the receptor (*e.g.*, A1) and those which were inert (*e.g.*, A28) (Fig. 2D). Collectively, these data demonstrate that highly specific, fully hIgG1 mAb were raised against LILRB3, amenable for the comprehensive evaluation of LILRB3 function.

### LILRB3 ligation modulates T cell activation and proliferation

Accordingly, using a select number of mAb, we sought to investigate the immunomodulating effect of the LILRB3 mAb on cellular effector functions. LILRB1 has previously been shown to directly inhibit T cell responses by causing dephosphorylation of the CD3 signaling cascade, and, in addition, has the potential to negatively regulate T cell activation by competing with CD8 for HLA-I binding (*31, 32*). Moreover, LILRBs can indirectly inhibit T cell responses by rendering antigen-presenting cells (APC), such as monocytes and DCs tolerogenic (*14, 18, 33*). To investigate the immunomodulatory potential of LILRB3 and its ability to regulate adaptive immune responses, we utilized a T cell proliferation assay incorporating fresh PBMCs isolated from healthy human donors, as before (*34*). Fcγ receptors (FcγRs) help mediate the effects of human IgG (*35*), therefore, to study the direct F(ab):LILRB3-mediated effects of the mAb on T cell proliferation, they were first deglycosylated to reduce FcγR-IgG interactions. SDS-PAGE confirmed a decrease in molecular weight of deglycosylated mAb compared to WT controls, indicative of successful deglycosylation (Fig. 3A). The mAb were then introduced to a T cell proliferation assay where CD3 and CD28 antibodies elicit cell clustering and CFSE dilution, indicative of a significant increase in CD8^+^ T cell proliferation, compared to non-treated controls (Fig. 3B and C). Clone A1, shown to be an agonist (Fig. 2D), significantly inhibited CD8^+^ T cell proliferation in this assay when compared to the isotype control (Fig. 3B and C). Other LILRB3-specific mAb had either no, or subtle effects, as represented by clones A16 and A28. These data demonstrate that LILRB3 ligation by agonistic mAb suppresses T cell responses; whereas, other clones confer no inhibitory effects. Similar effects were also observed when considering CD3^+^ CD8^−^ T cells (predominantly CD3^+^ CD4^+^ T cells; Fig. 3C and not shown). When the assay was repeated with isolated T cells, no inhibition was seen, confirming that APCs within the PBMC mixture, most likely monocytes, were responsible for the effects observed (fig. S3), as expected, given the lack of expression of LILRB3 on T cells (Fig. 1F and G).

**Fig. 3.**
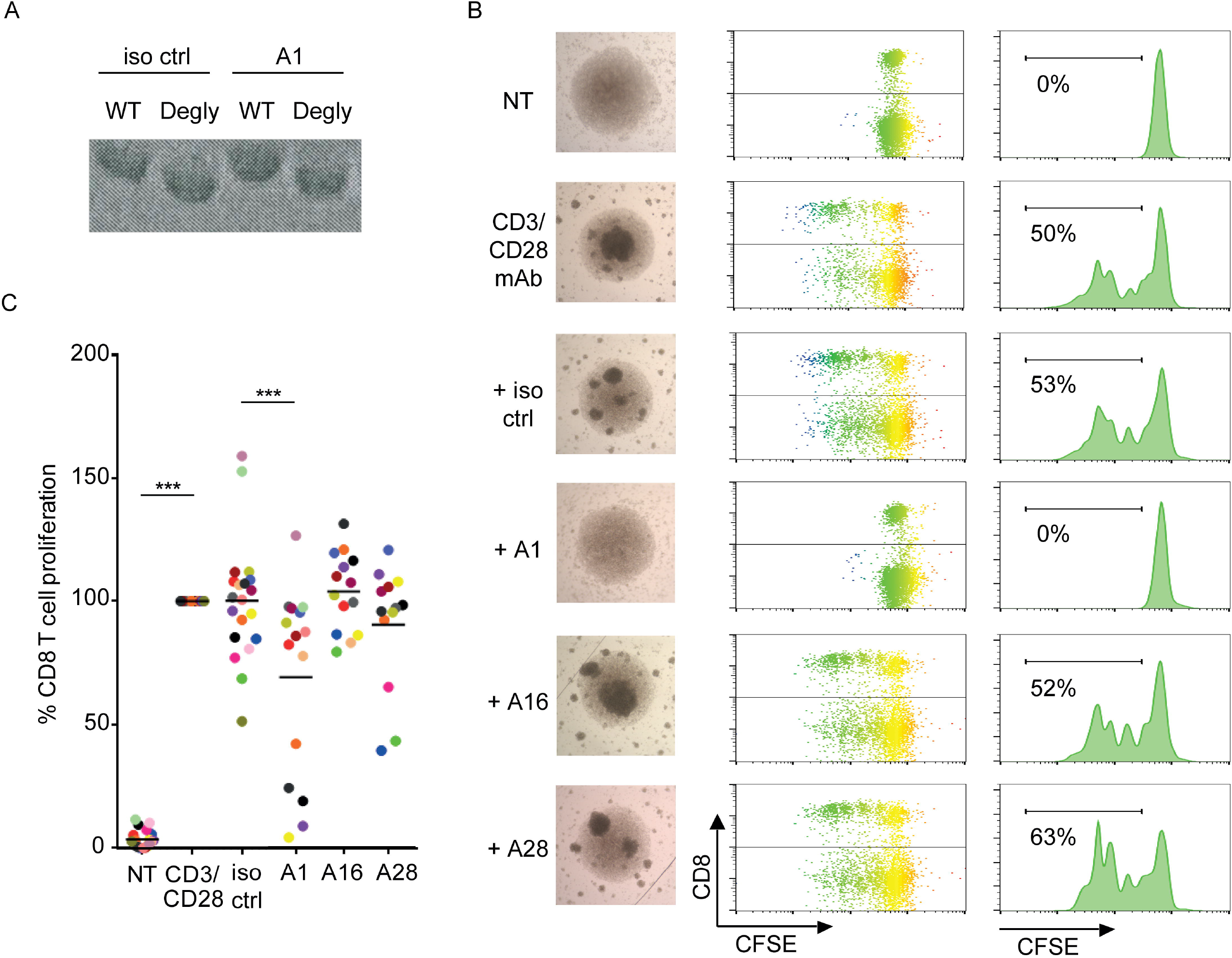
LILRB3 ligation regulates T cell activation and proliferation. CFSE-labelled PBMCs were stimulated with antibodies against human CD3 (0.02 μg/ml) and CD28 (5 μg/ml) in the presence or absence of isotype control (iso ctrl) or LILRB3 mAb (10 μg/ml) and proliferation measured through CFSE dilution after 3-5 days. (**A**) LILRB3 mAb were deglycosylated (Degly) through PNGase-treatment, as confirmed by SDS-PAGE; representative clone A1 shown. (**B**) Assessing T cell activation and proliferation following treatment. Light microscopy images following PBMC stimulation in culture. CD8^+^ T cell proliferation was assessed through CFSE dilution; representative plots, histograms (% proliferation indicated) and microscopy images shown (10x magnification). (**C**) Assessing the effects of deglycosylated LILRB3 mAb on T cell proliferation. CFSE dilution of CD8^+^ T cells, treated with the representative LILRB3 mAb was assessed by flow cytometry. Data normalized to anti-CD3/CD28-treated samples and mean represented by solid bars. Two-tailed paired T-test performed (*** *p* < 0.0001); *n* = 13-20 independent donors (each color represents an individual donor).

### LILRB3 ligation induces immune tolerance in humanized mice

Given these data showing that T cells could be suppressed following LILRB3 ligation on myeloid cells, we next investigated the possible effects of LILRB3 modulation in an allogeneic engraftment model using humanized mice, previously reconstituted with primary human fetal hematopoietic stem/progenitor cells (HSPC) (Fig. 4A). Characterization of peripheral blood leukocytes and bone marrow of adult humanized mice demonstrated that LILRB3 was expressed on, and restricted to, myeloid cells, but not lymphocytes, similar to humans (Fig. 4B and fig. S4). We recently showed that allogeneic human lymphoma cells are readily rejected in humanized mice due to HLA mismatch (*36*). To test the potential of LILRB3 ligation to suppress the allogeneic immune response, adult humanized mice were treated with the agonistic LILRB3 mAb (A1) and the engraftment of allogeneic human B cell lymphoma cells, derived from an unrelated donor (*36, 37*), monitored overtime (Fig. 4A). LILRB3 mAb treatment was able to induce a state of tolerance *in vivo* and led to a successful engraftment of the donor allogeneic cells (Fig. 4C). Accordingly, LILRB3-treated tumor-bearing humanized mice subsequently succumbed to disease with high tumor burden, whereas, isotype control-treated mice readily rejected the lymphoma cells without morbidity (Fig. 4D). These observations corroborate our *in vitro* functional assays and identify LILRB3 as a key regulator of immune tolerance in an allotransplant setting. Given the expression pattern of LILRB3 on myeloid but not lymphocytic cells in both the human PBMC and humanized mice, we sought to explore the effects of the LILRB3 mAb on these cells.

**Fig. 4.**
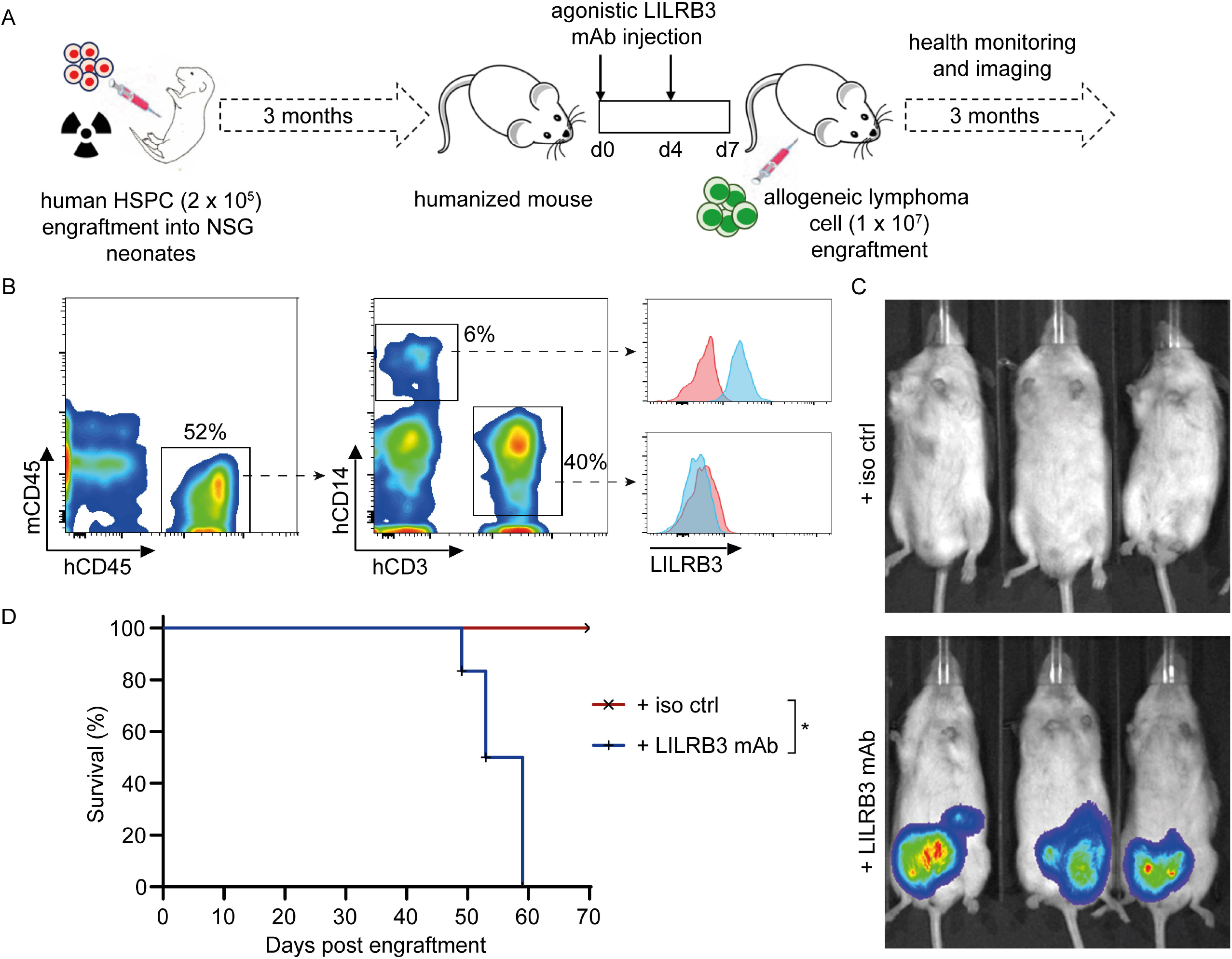
LILRB3 ligation induces tolerance *in vivo*. (**A**) Schematic of the generation of humanized mice and subsequent treatment regimens and monitoring. (**B**) Expression of LILRB3 on human myeloid cells in humanized mice. Representative flow cytometry plots (gated on live single cells) showing gating strategy and the restricted expression of LILRB3 on hCD45^+^ peripheral blood hCD14^+^ myeloid cells; isotype control in pink, LILRB3 in blue. (**C**) The effect of agonistic LILRB3 mAb on engraftment of allogeneic cells in humanized mice. Age- and sex-matched humanized mice were injected with 200 μg LILRB3 mAb (clone A1) or an isotype-matched (hIgG1) control mAb (iso ctrl) on day 0 and 4, i.v. and i.p, respectively. On day 7, mice were injected i.p. with 1×10^7^ non-autologous luciferase^+^ human lymphoma cells. Lymphoma cell growth was monitored over time using an IVIS imager; and (**D**) humanized mice were sacrificed upon the development of signs of terminal tumor development. Survival data was analyzed using log-rank test (*p* < 0.01); representative data from 3 independent experiments (3 individual HSPC donors) shown (*n* = 3 mice/group).

### LILRB3 ligation leads to transcriptional modification and M2-skewing of human APCs

To investigate the pathways and factors involved in LILRB3-mediated immunosuppression, we next investigated the transcriptomic changes in monocytes following LILRB3 engagement. Short-term (~18 hour) *in vitro* treatment of freshly-isolated human peripheral CD14^+^ monocytes with the agonistic LILRB3 mAb (A1) caused a dramatic shift in their phenotype (Fig. 5A), with the cells displaying an elongated morphology resembling immunosuppressed M2 macrophages (*38*). In accordance with this, RNAseq analysis revealed that ligation of LILRB3 on monocytes induced a signature resembling ‘M2-skewed’ immunosuppressive macrophages (Fig. 5B). Concurrently, the expression of genes associated with ‘M1-skewed’ immunostimulatory macrophages was downregulated in LILRB3-ligated monocytes (Fig. 5B-C). These data were confirmed by qPCR for a number of the differentially regulated genes on a further 6 donors (Fig. 5D). As further evidence, we showed that the effects were dependent upon LILRB3 agonism as treatment of monocytes with a non-agonistic LILRB3 mAb (A28), despite binding the same domain, did not affect monocyte phenotype or gene expression (Fig. 5D). Gene set enrichment analysis (GSEA) of the RNAseq data showed a positive correlation with gene signatures reported for suppressive macrophages, *e.g.*, oxidative phosphorylation (*39*). Conversely, LILRB3-ligated monocyte gene signatures negatively correlated with those reported for inflammatory macrophages, *e.g.*, IFN-γ and IFN-α responsive elements, as well as allograft rejection (Fig. 5E), in-line with our *in vivo* observations (Fig. 4). In summary, these data confirm that LILRB3 activation results in significant phenotypic and transcriptional alterations in human primary myeloid cells, leading to potent inhibition of downstream immune responses.

**Fig. 5.**
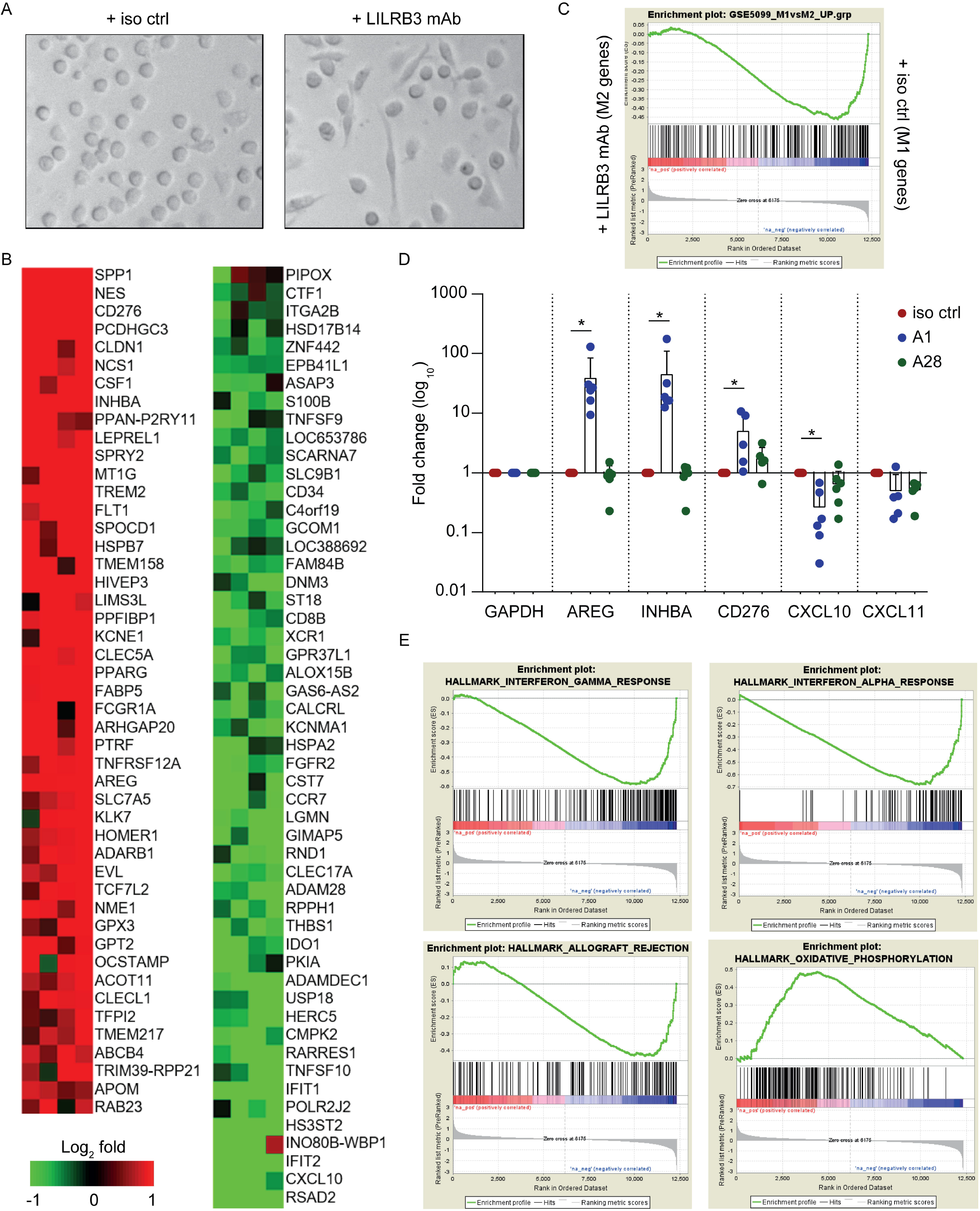
Human LILRB3 ligation reprograms human primary myeloid cells. Freshly isolated human peripheral CD14^+^ monocytes were treated with an isotype control (iso ctrl) or a human LILRB3 mAb (clone A1) and then assessed. (**A**) Agonistic LILRB3 mAb affects monocyte morphology. Light microscopy images following overnight treatment of freshly-isolated CD14^+^ monocytes with indicated mAb in culture (10× magnification). (**B**) Transcriptomic analysis of LILRB3-treated monocytes reveals upregulation of M2-associated genes compered to controls. RNA was extracted from cells following mAb treatment (~18 hours) and subjected to RNA sequencing. Red depicts genes that were significantly upregulated and green depicts genes that were significantly downregulated compared to isotype control treated-cells (*n* = 4 independent donors). (**C**) Ligation of LILRB3 on primary human CD14^+^ monocytes induces M2-polarized genes. GSEA graph showing a significant enrichment for M2-polarizing genes in LILRB3-treated monocytes versus isotype control, respectively. UP; upregulated, NES; normalized enrichment score = −1.68; FWER; familywise-error rate *p* <0.001. (**D**) qPCR analysis of selected genes following LILRB3 ligation on monocytes using an agonistic LILRB3 mAb (A1), a non-agonistic LILRB3 mAb (A28) or an isotype control (iso). Data were normalized to GAPDH mRNA levels and standardized to the levels of isotype control-treated monocytes. Fold difference data were log_10_ transformed. One-way ANOVA with Bonferroni’s multiple comparisons test was performed, *n* = 5-6 independent donors (* *p* < 0.005). (**E**) GSEA analysis showing negative correlation with ‘IFN-γ’ (NES=-2.17; FWER *p* < 0.001), ‘IFN-α’ (NES=-2.3; FWER *p* < 0.001) and ‘allograft rejection’ (NES=-1.58; FWER p = 0.14) signaling elements and positive correlation with ‘oxidative phosphorylation’ (NES=2; FWER *p* < 0.001).

## Discussion

We previously demonstrated that ligation of LILRB1 on human DCs induces a tolerogenic phenotype, hindering T cell responses (*18, 40*). In this study, we investigated another inhibitory LILR family member, LILRB3, whose function, largely due to lack of suitable reagents and experimental systems, is not yet fully determined. Limited previous studies investigated the consequences of LILRB3 activation on granulocytes and have demonstrated its inhibitory function on neutrophils (*41*) and basophils (*42*) in culture. Here, we largely concentrated on myelomonocytic cells and the subsequent regulation of adaptive immune responses. We, therefore, initially generated and characterized an extensive panel of fully human mAb with specificity for LILRB3 through a number of stringent panning and selection processes. Those clones showing cross-reactivity to other human LILR family members were excluded. Immunoprofiling of circulatory leukocytes from healthy donors using these highly specific mAb confirmed the reported expression of LILRB3 on myelomonocytic and granulocytic cells, but not on lymphocytes (*2, 3, 6*). This pattern of expression on myeloid and granulocytic but not lymphoid cells was confirmed in a large cohort of independent donors (>50), suggesting that, despite the polymorphic nature of LILRB3 (*2, 9, 43*), the selected antibodies recognize many, if not all, variants, which is important for the development of these reagents for therapeutic applications. Subsequent analysis showed that the LILRB3 mAb displayed a range of affinities, albeit all within the nanomolar (nM) range, with similar on-rates, but off-rates differing over three orders of magnitude. K_*D*_ values in the low nM range are generally considered to be viable drug candidates; rituximab, for example, has an 8 nM affinity for its target, CD20 (*44*). This suggests that the LILRB3 mAb generated here have potential as therapeutic agents. However, as LILRB3 shares high sequence homology (>95%) in its extracellular domain with LILRA6, there is a possibility that some LILRB3 mAb may also recognize shared epitopes on LILRA6, if co-expressed (*9*). These initial data might receive further evidence from other reagents as well as investigation as to whether LILRA6 protein is detectable in leukocyte subsets, *e.g.*, using proteomics approaches similar to a recent study with neutrophils (*41*). Epitope mapping experiments revealed that the specific LILRB3 mAb reported here were generated against two specific ectodomains, either Ig-like domain two or four. Interestingly, none of the generated specific LILRB3 mAb bound to Ig-like domains one or three, suggesting that these domains may not contain epitopes that are unique for LILRB3, or more likely those unique residues are not exposed/accessible. Collectively, these data confirm that our LILRB3 mAb will be useful tools for dissecting LILRB3’s molecular mechanisms and may additionally have therapeutic benefits in relevant pathologies.

The ability of the LILRB3 mAb to influence T cell responses was variable: ranging from inhibition to indications of modest increases in proliferation, supportive of agonistic or blocking properties, respectively. Similar to LILRB1 (*17, 18, 21*), these effects are likely through manipulations of APCs, specifically monocytes, as they are the only cells expressing LILRB3 in the culture. In support of this, the agonistic LILRB3 mAb did not suppress T cell proliferation in the absence of monocytes. Binding epitopes influence the ability of mAb to modulate receptor function in many systems (*35, 45*) and so it was unsurprising to see LILRB3 mAb capable of differing functions. However, the D4-binding A1 mAb was a strong inhibitor of proliferation; whereas, other D4-binding mAb (*e.g.*, A28) had no significant effect. Therefore, domain-specific epitopes did not seem to correlate directly with LILRB3 mAb-mediated effector cell functions and may not be predictive of LILRB3 mAb function *per se*. Further detailed analyses, *e.g.*, surface alanine scanning mutagenesis (*45*) and/or structural studies, are required to define the specific extracellular epitopes engaged by the selected LILRB3 mAb and to investigate their influence on receptor activity.

Our observations demonstrating immunoinhibitory activities downstream of LILRB3 were further confirmed in the reconstituted humanized mouse model. In this system, where LILRB3 is present only on the hematopoietic cells, and predominantly monocytes, in the absence of appreciable numbers of neutrophils, ligation of LILRB3 with an agonistic LILRB3 mAb prior to injection of allogeneic lymphoma cells (*36, 37*) induced tolerance *in vivo* and enabled subsequent tumor engraftment. This demonstrates the capacity of LILRB3 to exert profound immunosuppressive effects that may be exploited in therapeutic settings, such as autoimmunity and transplantation, where transient induction of immune tolerance will be beneficial. Similar observations were previously reported using a LILRB1 transgenic mouse model, where interactions between LILRB1 and MHCI or HLA-G expanded MDSCs and prolonged allogenic graft survival *in vivo* (*46, 47*).

Although typically regarded as an orphan receptor, earlier studies suggest that LILRB3 may associate with cytokeratin-associated proteins such as those exposed on necrotic cancer cells (*30*). Others have also identified angiopoietin-like protein 5 and bacteria, such as *Staphylococcus aureus*, as a source of potential ligands (*48, 49*). Therefore, our data provide a strong mechanism of action whereby such endogenous or pathogenic ligands may be able to subvert immune responses by ligating LILRB3 during an ongoing immune response.

To investigate the pathways and factors involved in LILRB3-mediated immunosuppression, we investigated the transcriptomic changes in isolated peripheral myeloid cells following LILRB3 activation. Over one hundred genes were differentially regulated in primary human monocytes following LILRB3 ligation, some of which are known to be modulated in M2 macrophages (*13, 50*). Amphiregulin (AREG) was among the genes whose expression was significantly upregulated in LILRB3-ligated monocytes. AREG is an epidermal growth factor-like growth factor, responsible for inducing tolerance and immunosuppression, via various mechanisms including enhancement of Treg activity (*51*). Furthermore, AREG is overexpressed in tumor-associated DCs (*52*) and suppressive/M2 macrophages (*53*) and has been suggested to play a crucial role in immunosuppression and cancer progression (*54*). Such LILRB3-inducible factors may be responsible for the suppression observed in our T cell assays. Our ongoing efforts aim to interrogate these findings further and define the mechanisms responsible for LILRB3-mediated suppression of immune responses at molecular and cellular levels, *e.g.*, via siRNA knockdown of AREG in monocytes and validation of differentially regulated genes in the humanized mouse models. A recent study investigating the mode of action of Glatiramer acetate (Copaxone), a peptide-based drug licensed in the late 1990’s, used to treat patients with the relapsing-remitting form of multiple sclerosis that ameliorates autoimmunity, identified LILRB2 and LILRB3 as potential ligands (*55*). On the other hand, blocking human LILRB2 with antagonistic mAb on human myeloid cells is able to promote their pro-inflammatory activity and enhance antitumor responses in preclinical models (*56*); and a LILRB2 mAb (MK-4830) recently entered phase I clinical trials (NCT03564691) for advanced solid tumors. Furthermore, recent data by Zhang and colleagues suggest that LILRB4 signaling in leukemia cells mediates T cell suppression and supports tumor cell dissemination to distal organs (*57*). These recent compelling reports further support our findings, demonstrating that activation of human LILRB3 induces immunosuppression via reprogramming of myeloid cells (*i.e*., reducing M1-like maturation and promoting suppressive function).

In conclusion, our findings show that LILRB3 engagement on primary human myeloid cells exerts potent immunoinhibitory functions and that LILRB3-specific mAb are potentially powerful immunomodulatory agents, with broad application ranging from transplantation to autoimmunity and beyond, where fine-tuning of immune responses through myeloid cell activity is desired.

## Materials and Methods

### Ethics Statement

All research with human samples and mice was performed in compliance with institutional guidelines, the Declaration of Helsinki and the US Department of Health and Human Services Guide for the Care and Use of Laboratory Animals. The Committee on Animal Care at Massachusetts Institute of Technology (MIT) reviewed and approved the studies described here. All human samples (adult peripheral blood and fetal liver) were collected anonymously with informed consent by a third party and purchased for research. For human peripheral blood, ethical approval for the use of clinical samples was obtained by the Southampton University Hospitals NHS Trust; from the Southampton and South West Hampshire Research Ethics Committee following provision of informed consent. Primary chronic lymphocytic leukemia (CLL) samples were released from the Human Tissue Authority licensed University of Southampton, Cancer Science Unit Tissue Bank as part of the LPD study REC number 228/02/T.

### Cell culture

Cell lines were grown at 37°C in either RPMI 1640 medium supplemented with 10% heat-inactivated fetal calf serum (FCS) (Sigma-Aldrich, UK), 100 U/ml Penicillin-Streptomycin, 2 mM Glutamine and 1 mM Pyruvate (Thermo Fisher Scientific, UK) in a humidified incubator with 5% CO_2_, Freestyle 293F media, in 8% CO_2_, shaking at 130 rpm, or Freestyle CHO media (Thermo Fisher Scientific, UK) with 8 mM Glutamine, in 8% CO_2_, shaking at 140 rpm.

### Antibody generation and production

#### Generation of LILRB3 antibodies

Generation of LILRB3-specific mAb was performed using the nCoDeR phage-display library (*28*). Three consecutive panning rounds were performed, as well as a pre-panning step. In the panning, human (h) Fc fusion proteins containing the extracellular domains of LILRB1 and LILRB3 (LILRB-hFc) were used as ‘non-target’ or ‘target’, respectively. These proteins were produced in transiently transfected HEK293 cells followed by purification on protein A, as described previously (*35*). CHO-S cells transiently transfected to express the various LILRB proteins were also used as targets/non-targets in the panning.

In panning 1, BioInvent n-CoDeR® scFv were selected using biotinylated in-house produced recombinant LILRB3-hFc recombinant fusion proteins (captured with streptavidin-coated Dynabeads®) with or without competition or LILRB1-hFc coated to etched polystyrene balls (Polysciences, US) or plastic immunotubes. Binding phages were eluted by trypsin digestion and amplified on plates using standard procedures (*58*). The amplified phages from panning 1 were used for panning 2, the process repeated, and the amplified phages from panning 2 used in panning 3. In the third panning round, however, amplified phages from all 3 strategies were combined and selected against LILRB–expressing CHO-S cells, prior to making the final LILRB3-specific mAb selection.

Next, the LILRB3-positive scFv from the enriched phage repertoires from panning 3 were recloned to allow soluble scFv expression in *E. coli*. The soluble scFv fragments expressed by individual clones were tested for binding against LILRB-transfected CHO-S cells using FMAT, and recombinant LILRB protein by ELISA. This allowed the identification of clones binding specifically to LILRB3. Clones were then further reduced in a tertiary screen against CHO-S cells expressing LILRB1-3 and primary cells (PBMCs) using a high throughput flow cytometry screening system and data analyzed by TIBCO Spotfire® software (TIBCO, USA). Clones showing specific patterns of binding to LILRB3 were sequenced, yielding LILRB3-specific mAb.

#### Production of full-length IgG

The unique scFv identified above were cloned into a eukaryotic expression system allowing transient expression of full-length IgG in HEK293-EBNA cells. The antibodies were then purified from the culture supernatants using Protein A affinity chromatography as previously described (*35*).

#### Production of deglycosylated IgG

To allow dissection of Fc- and Fab-dependent effector functions, IgG were deglycosylated using PNGase F (Promega) with 0.05 U of PNGase/μg of IgG, at 37°C for at least 15 hours. Deglycosylation was confirmed by reduction in size of the heavy chain on SDS-PAGE.

### Production of domain mutant constructs

Using wild-type LILRB3 cDNA isolated from a healthy donor PBMCs, a series of domain mutant DNA constructs were generated by overlap PCR to express 1, 2 or 3 LILRB3 Ig-like domains (with domains identified based on annotations in Uniprot) for comparison to WT LILRB3 (4 domains). The gene constructs were then cloned into pcDNA3.

### Cell Transfections

10 × 10^6^ HEK293F cells were transiently transfected with 10 μg of plasmid DNA by lipofection using 233 fectin with Optimem 1 Media (Thermo Fisher Scientific, UK).

### Preparation of human leukocytes

PBMC were isolated from leukocyte blood cones (Blood Transfusion Services, Southampton General Hospital) by gradient density centrifugation using lymphoprep (Axis-Shield, UK) and used for subsequent experiments, as before (*59*).

### Flow cytometry

For cell surface staining of PBMCs or whole blood, cells were blocked with 2% human AB serum (Sigma-Aldrich, UK) for 10 minutes on ice and then stained with the relevant APC-labelled mAb or hIgG1 isotype (BioInvent, Sweden), alongside the following cell surface markers: CD14-PE (eBioscience, UK), CD20-A488 (Rituximab, in house), CD3-PE-Cy7, CD56-APC-Cy7 or CD15-Pacific Blue, CD15-PE and CD66B-FITC (all Biolegend, UK). Cells were stained for 30 minutes at 4°C and then washed twice, first in 10% red blood cell (RBC) lysis buffer (Serotec, UK) for PBMCs or 1× Erythrolyse RBC Lysing Buffer (Biorad, UK) for whole blood, and then with FACS wash (PBS, 1% BSA, 10 mM NaN_3_), before acquisition on a FACSCalibur or FACSCanto II (BD Biosciences, USA) and analyzed with FlowJo software (BD Biosciences, USA).

For assays to determine if mAb bound to similar cross-blocking epitopes 1 × 10^6^ PBMCs were blocked with 2% human AB serum for 10 minutes and stained with 10 μg/ml unconjugated LILRB3 mAb for 30 minutes at 4°C. The cells were then stained with directly-conjugated LILRB3 mAb (clone 222821; mouse IgG2a; R&D Systems, UK) for 20 minutes at 4°C, before washing and acquisition using a FACSCalibur.

For LILRB3 epitope mapping studies, LILRB3-domain mutant-transfected HEK293F cells were stained with the relevant LILRB3 mAb for 25 minutes at 4°C, washed twice, stained with an anti-human-PE secondary (Jackson ImmunoResearch, USA) for 20 minutes at 4°C, before washing and acquisition using a FACSCalibur.

For staining of 2B4 reporter cells expressing LILR-A1, −A2, −A5, −B1, −B2, −B3, −B4, or −B5 (or non-transfected controls) (*30, 60*) cells were stained with 10 μg/ml LILRB mAb and incubated at 37°C with 5% CO_2_, overnight. The following day, the cells were washed and stained with a secondary anti-hIgG antibody (Jackson ImmunoResearch, USA) at 4°C, for 45 minutes. The cells were washed and acquisition performed using a FACScan (BD Biosciences, USA) and data analyzed with FlowJo software (BD Biosciences, USA).

### Surface plasmon resonance (SPR)

SPR was performed with the Biacore T100 (GE Healthcare, UK) as per the manufacturer’s instructions. LILRB3-hFc recombinant protein (extracellular LILRB3 domain with a hFc tag) was used as the ligand and immobilized by amine coupling onto a series S sensor chip (CM5). Various LILRB3 mAb were used as ‘analytes’ and flowed across the chip, and SPR measured. *K*_D_ values were calculated from the ‘Univalent’ model of 1:1 binding by Kd [1/s] / Ka [1/Ms], using the Biacore™ T100 Evaluation Software (GE Healthcare, UK).

### T cell Proliferation assay

PBMCs (1-2 × 10^7^) were labelled with 2 μM carboxyfluorescein succinimidyl ester (CSFE) at room temperature for 10 minutes. An equal volume of FCS was then added to quench labeling for 1 minute, prior to washing. Cells were subsequently resuspended in serum-free CTL medium (Immunospot, Germany) and plated at 1×10^5^ cells/well in a 96-well round-bottom plate (Corning, UK). Cells were then stimulated with 0.02 μg/ml CD3 (clone OKT3, in-house), 5 μg/ml CD28 (clone CD28.2; Biolegend, UK) and 10 μg/ml LILRB3 antibodies or a relevant isotype. Plates were then incubated at 37°C for 4 days, after which time cells were stained with 5 μg/ml CD8-APC (clone SK1; Biolegend, UK), harvested and CSFE dilution measured by flow cytometry, as a readout for T cell proliferation.

### Hematopoietic stem/progenitor cell (HSPC) isolation and generation of humanized mice

Humanized mice were generated, as described (*36*). In brief, human fetal livers were obtained from aborted fetuses at 15–23 weeks of gestation, in accordance with the institutional ethical guidelines (Advanced Bioscience Resources, Inc., USA). All women gave written informed consent for the donation of their fetal tissue for research. Fresh tissue was initially cut into small pieces and digested with collagenase VI (2 mg/ml; Roche) for 30 minutes at 37°C. Single-cell suspensions were prepared by passing the digested tissue through a 100 μm cell strainer (BD Biosciences, USA). HSPCs were purified using a CD34^+^ selection kit (Stem Cell Technologies, Canada); the purity of CD34^+^ cells was 90%–99%. Viability was determined through trypan blue exclusion of dead cells. All cells were isolated under sterile conditions and injected into NSG mice.

NSG mice were purchased from the Jackson Laboratories (Bar Harbor, USA) and maintained under specific pathogen-free conditions in the animal facilities at MIT. To reconstitute mice, newborn pups (less than 2 days old) were irradiated with 100 cGy using a Gamma radiation source and injected intracardially with CD34^+^ cells (~2 × 10^5^ cells/recipient), as reported previously (*36*). Around 12 weeks later, human leukocyte cell reconstitution of PBMCs was determined by flow cytometry and calculated as follows: % human CD45^+^ cell / (% human CD45^+^ cell + % mouse CD45^+^ cell). Mice with ≥ 40% human CD45^+^ leukocytes were used in subsequent experiments.

### *In vivo* allograft assay

Fully reconstituted humanized mice were injected with 200 μg LILRB3 mAb (clone A1) or an isotype-matched (hIgG1) control on day 0 and day 4, i.v. and i.p, respectively. On day 7, cohorts of mice were injected i.p. with 1 × 10^7^ luciferase-positive human ‘double-hit’ B cell lymphoma cells (*36, 37*), derived from unrelated donors. Lymphoma cell growth was monitored over time using an IVIS Spectrum-bioluminescent imaging system, as before (*36*). Mice with palpable tumors were sacrificed and Kaplan-Meier survival curves plotted.

### Transcriptome analysis

To assess LILRB3-mediated transcriptional changes on monocytes, human peripheral blood monocytes were isolated from freshly prepared PBMCs taken from healthy donors using an EasySep™ Human Monocyte Enrichment Kit (negative selection cell; StemCell Technologies, Canada). Cells were incubated in CTL medium supplemented with 100 U/ml Penicillin-Streptomycin, 2 mM Glutamine and HEPES buffer and treated with 10 μg/ml of an isotype control or an agonistic LILRB3 mAb (clone A1; hIgG1). 18 hours later, cells were lysed in RLT lysis buffer containing β-mercaptoethanol and total RNA extracted using the RNeasy micro kit (Qiagen, USA). Total RNA was assessed for quality and quantified using a total RNA 6000 Nano LabChip on a 2100 Bioanalyzer (Agilent Inc., USA) and cDNA libraries prepared and sequenced according to the Illumina TruSeq RNA Sample Preparation Guide for SMARTer Universal Low Input RNA Kit (Clontech, USA) and a HiSeq 2000 system (Illumina, USA). RNAseq outputs were aligned to hg19 using Bowtie2 v2.2.3 (*61*). The number of mapped reads were quantified by RSEM v1.2.15 (*62*). Differential expression analysis between paired samples before and after treatment was performed using edgeR (*63*) with *p* <0.05 and >2 fold-change cut-offs. Differentially expressed genes were annotated using online functional enrichment analysis tool DAVID (http://david.ncifcrf.gov/) (*64*). Gene-set enrichment analysis (GSEA) was performed using Broad Institute Software (*65*), with the gene list pre-ranked according to logFC values from the edgeR output. For comparison of gene-set expression, M1 and M2 macrophage gene sets (*50*) were obtained from the Molecular Signature Database (http://software.broadinstitute.org/gsea/msigdb/). Heatmaps were visualized with MeV (*66*). Raw sequences are deposited in the database of Gene Expression Omnibus with accession ID GSE151675.

### Quantitative PCR (qPCR)

Probe-based qPCR was used to amplify cDNA in 20μl reactions performed in triplicate for each sample condition in a PCR plate (Bio-Rad, UK), as per the manufacturer’s protocol. The 96-well plate was run on a C1000 Thermal Cycler CFX96 Real-time System PCR machine (Bio-Rad, Kidlington, UK). The CFX manager software (Bio-Rad, Kidlington, UK), was used for data acquisition and analysis of gene expression initially recorded as cycle threshold values (Ct). The Ct values were normalised to housekeeping gene Glyceraldehyde 3-phosphate dehydrogenase (GAPDH) and standardised to gene expression levels in isotype control-treated samples.

### Statistics

Paired two-tailed T-tests were performed for both the immunophenotyping and T cell proliferation data; straight bars indicate median values. On bar graphs, where at least 3 experiments were performed, error bars represent standard deviation. Kaplan-Meier plots were analyzed by Log-rank test. One-way ANOVA with Bonferroni’s multiple comparisons test were performed for qPCR data analysis. Statistical analysis was performed using GraphPadPrism (v6-8).

## Authorship

Contribution: AR and MSC were grant holders, initiated the research proposals, supervised the project and wrote the manuscript. MY, TS, UM, UT, BH, AL and MM generated human LILRB3 mAb and performed the initial screening. MY characterized the mAb and performed the *in vitro* functional assays. HTCC supported the molecular biology and CP and JSM supported the *in vitro* assays. JT provided reagents and edited the manuscript. BF provided expert advice on the mAb generation and edited the manuscript. JC provided expert advice on the *in vivo* assays and edited the manuscript. AR performed the *in vivo* experiments in humanized mice. DCJ, designed DNA constructs used to generate antibodies, performed specificity assays against LILR transfectants and edited the manuscript. CIM performed SPR assays. GH and SMT analyzed the RNAseq data. IT and MJG provided expert advice on the epitope mapping and functional assays, respectively, and edited the manuscript.

## Acknowledgments and Grant Support

We would like to thank the Antibody and Vaccine Group antibody production team, Profs Tim Elliott (University of Southampton) and Jean-Michel Sallenave (Université Paris Diderot) for review of the manuscript; Drs Francesco Forconi and Ian Tracey (University of Southampton) for provision of CLL samples. We would like to thank the Koch Institute Flow Cytometry (Glenn Paradis), Bioinformatics and Computing (Stuart Levin) and Animal Imaging and Preclinical Testing cores (Scott Malstrom) for their technical assistance.

This work was supported by an iCASE studentship awarded to AR, BF and MSC to support MY from the BBSRC and BioInvent International AB, Sweden. AR was a recipient of a Blood Cancer UK Visiting Fellowship (14043). Work in the JT lab was supported by European Research Council (ERC) under the European Union’s Horizon 2020 research and innovation programme (grant agreement No: 695551).

## Supplementary figure legends

**Fig. S1.**
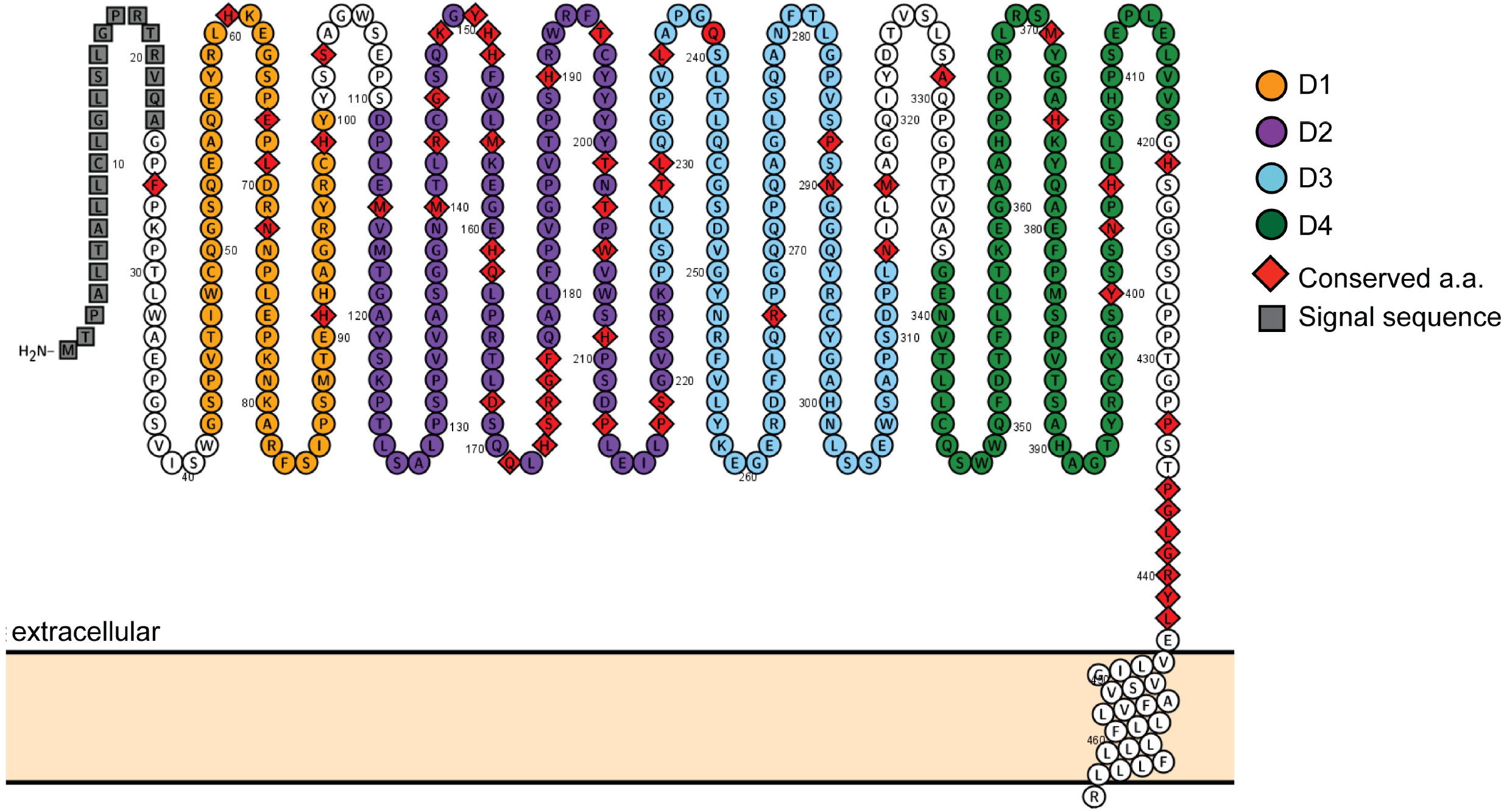
Topological structure of LILRB3 ectodomain. Topological representation of the predicted extracellular domains of human LILRB3 (accession no. O75022) and its conserved a.a. residues, as aligned against LILRB1 (accession no. Q8NHL6), LILRB2 (accession no. Q8N423), LILRB4 (accession no. Q8NHJ6) and LILRB5 (accession no. O75023). Each predicted Ig-like domain (D1-4) has been separately color-coded and the conserved a.a. residues identified by red rhombus symbols. Source: UniProt and Protter.

**Fig. S2.**
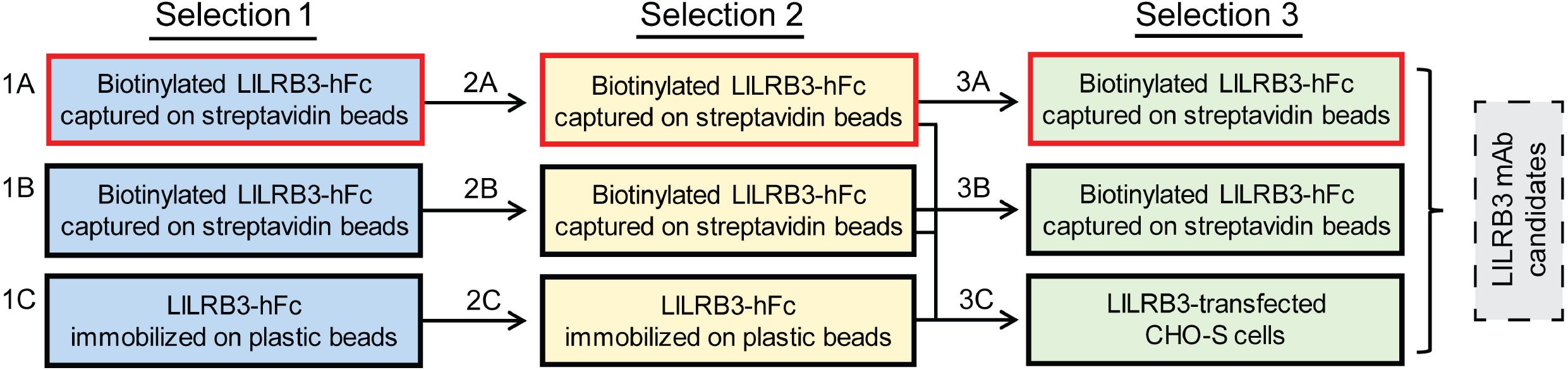
Schematic of LILRB3-specific mAb selection strategy. Three ‘selection’ rounds were performed to generate LILRB3-specific mAb by phage display. Three different methods to display the target protein were utilized: biotinylated target captured on streptavidin magnetic beads with competition *(A)*, biotinylated target without competition *(B)*, and immobilized protein coated on plastic polystyrene beads in the first two selection rounds, followed by cells expressing the target in selection around three using all phages from all three techniques used in panning two *(C)*. Throughout the panning rounds, LILRB1 was used as a non-target and as a competitor for LILRB3. 50 nM biotinylated non-target was used in the pre-selection for strategies where biotinylated protein was used *(A, B)*. 10 μg/ml non-target was used for strategies where coated protein was used *(C)*. No pre-selection was required for the cell strategies *(3C)*. Red border indicates competition with excess non-target protein was performed.

**Fig. S3.**
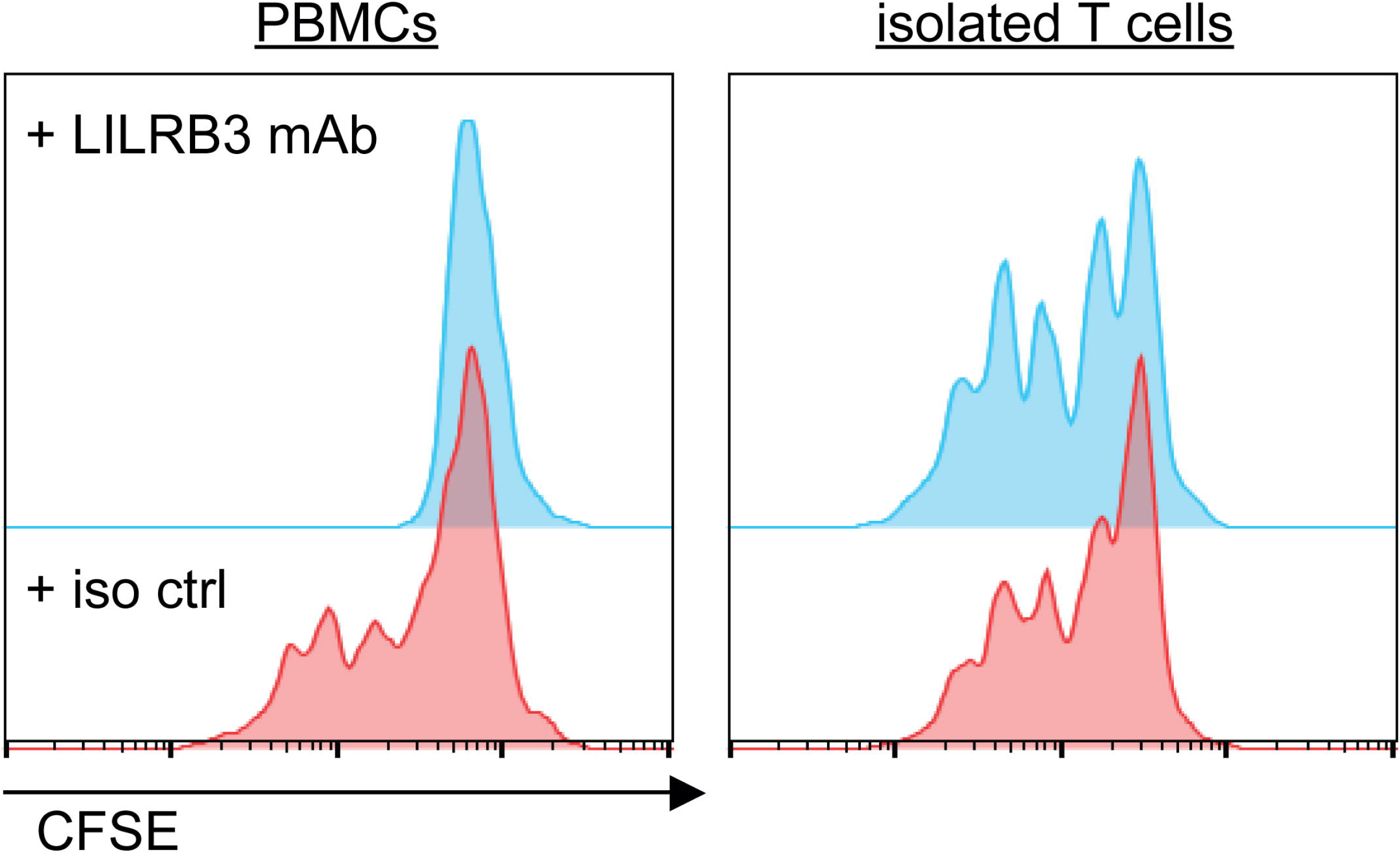
T cell proliferation is not affected by agonistic LILRB3 mAb in the absence of myeloid cells. Fresh PBMCs or purified T cells, isolated from matched PBMCs using a human CD3 negative selection kit, were CFSE-labeled and then cultured in the presence of soluble (0.02 μg/ml) or plate-bound (5 μg/ml) OKT3 mAb and either 10 μg/ml of isotype control (pink) or an agonistic LILRB3 mAb (blue). Proliferation was measured through CFSE dilution on day 4 by flow cytometry. Representative histogram of 3 independent experiments shown for each culture.

**Fig. S4.**
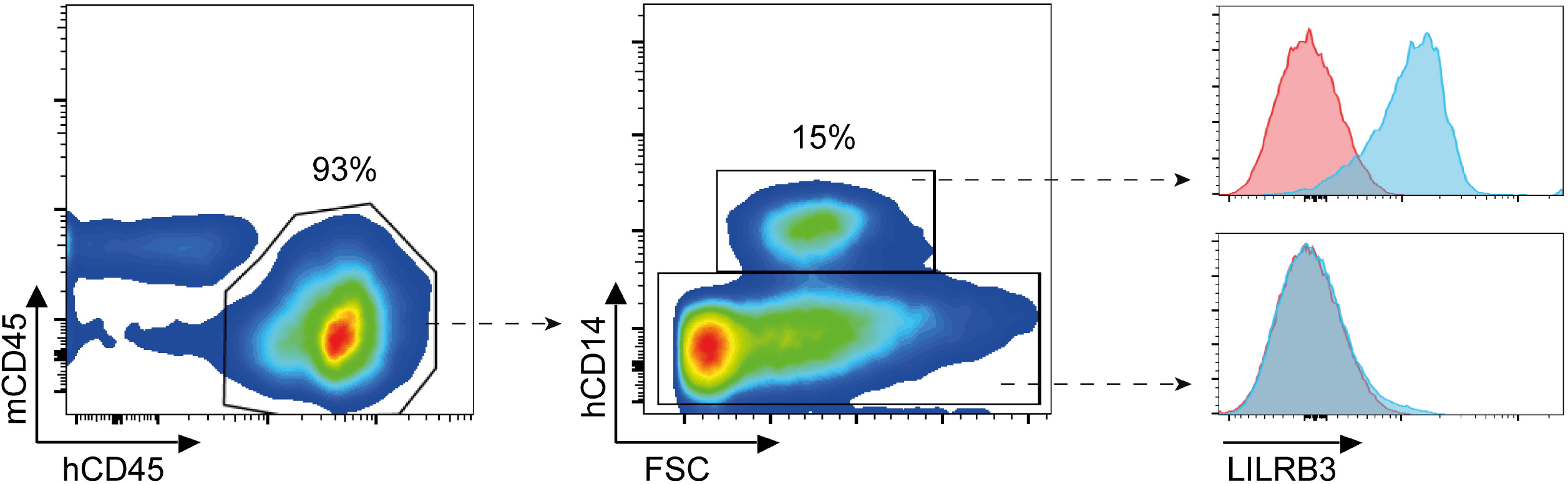
Expression of LILRB3 on bone marrow-resident myeloid cells in humanized mice. Expression of LILRB3 on freshly isolated hCD45^+^ bone marrow myeloid cells in humanized mice. Representative (3 mice/group, from 3 independent HSPC donors) flow cytometry plots (gated on live single cells) showing gating strategy and the restricted expression of LILRB3 on hCD14^+^ myeloid cells ~3 months post engraftment of HSPCs; isotype control in pink, LILRB3 in blue.

## References

1. L. Borges, M. L. Hsu, N. Fanger, M. Kubin, D. Cosman, A family of human lymphoid and myeloid Ig-like receptors, some of which bind to MHC class I molecules. J Immunol 159, 5192–5196 (1997).

2. M. Colonna et al., A common inhibitory receptor for major histocompatibility complex class I molecules on human lymphoid and myelomonocytic cells. The Journal of experimental medicine 186, 1809–1818 (1997).

3. W. van der Touw, H. M. Chen, P. Y. Pan, S. H. Chen, LILRB receptor-mediated regulation of myeloid cell maturation and function. Cancer Immunol Immunother 66, 1079–1087 (2017).

4. L. E. Hudson, R. L. Allen, Leukocyte Ig-Like Receptors - A Model for MHC Class I Disease Associations. Front Immunol 7, 281 (2016).

5. D. N. Burshtyn, C. Morcos, The Expanding Spectrum of Ligands for Leukocyte Ig-like Receptors. J Immunol 196, 947–955 (2016).

6. L. Borges, M. L. Hsu, N. Fanger, M. Kubin, D. Cosman, A family of human lymphoid and myeloid Ig-like receptors, some of which bind to MHC class I molecules. J Immunol 159, 5192–5196 (1997).

7. H. Nakajima, J. Samaridis, L. Angman, M. Colonna, Human myeloid cells express an activating ILT receptor (ILT1) that associates with Fc receptor gamma-chain. J Immunol 162, 5–8 (1999).

8. C. C. Chang et al., Polymorphism and linkage disequilibrium of immunoglobulin-like transcript 3 gene. Hum Immunol 69, 284–290 (2008).

9. M. R. Lopez-Alvarez, D. C. Jones, W. Jiang, J. A. Traherne, J. Trowsdale, Copy number and nucleotide variation of the LILR family of myelomonocytic cell activating and inhibitory receptors. Immunogenetics 66, 73–83 (2014).

10. K. Hirayasu, H. Arase, Functional and genetic diversity of leukocyte immunoglobulin-like receptor and implication for disease associations. J Hum Genet 60, 703–708 (2015).

11. F. W. Velten, K. Duperrier, J. Bohlender, P. Metharom, S. Goerdt, A gene signature of inhibitory MHC receptors identifies a BDCA3(+) subset of IL-10-induced dendritic cells with reduced allostimulatory capacity in vitro. Eur J Immunol 34, 2800–2811 (2004).

12. C. C. Chang et al., Tolerization of dendritic cells by T(S) cells: the crucial role of inhibitory receptors ILT3 and ILT4. Nat Immunol 3, 237–243 (2002).

13. M. Beyer et al., High-resolution transcriptome of human macrophages. PLoS One 7, e45466 (2012).

14. J. S. Manavalan et al., High expression of ILT3 and ILT4 is a general feature of tolerogenic dendritic cells. Transpl Immunol 11, 245–258 (2003).

15. J. Banchereau et al., Immunoglobulin-like transcript receptors on human dermal CD14+ dendritic cells act as a CD8-antagonist to control cytotoxic T cell priming. Proceedings of the National Academy of Sciences of the United States of America 109, 18885–18890 (2012).

16. N. A. Fanger et al., The MHC class I binding proteins LIR-1 and LIR-2 inhibit Fc receptor-mediated signaling in monocytes. Eur J Immunol 28, 3423–3434 (1998).

17. A. A. Barkal et al., Engagement of MHC class I by the inhibitory receptor LILRB1 suppresses macrophages and is a target of cancer immunotherapy. Nat Immunol 19, 76–84 (2018).

18. N. T. Young et al., The inhibitory receptor LILRB1 modulates the differentiation and regulatory potential of human dendritic cells. Blood 111, 3090–3096 (2008).

19. M. K. Rochat et al., Maternal vitamin D intake during pregnancy increases gene expression of ILT3 and ILT4 in cord blood. Clin Exp Allergy 40, 786–794 (2010).

20. M. Brenk et al., Tryptophan deprivation induces inhibitory receptors ILT3 and ILT4 on dendritic cells favoring the induction of human CD4+CD25+ Foxp3+ T regulatory cells. J Immunol 183, 145–154 (2009).

21. C. S. Wagner et al., Human cytomegalovirus-derived protein UL18 alters the phenotype and function of monocyte-derived dendritic cells. J Leukoc Biol 83, 56–63 (2008).

22. M. G. Petroff, P. Sedlmayr, D. Azzola, J. S. Hunt, Decidual macrophages are potentially susceptible to inhibition by class Ia and class Ib HLA molecules. J Reprod Immunol 56, 3–17 (2002).

23. R. Apps, L. Gardner, A. M. Sharkey, N. Holmes, A. Moffett, A homodimeric complex of HLA-G on normal trophoblast cells modulates antigen-presenting cells via LILRB1. Eur J Immunol 37, 1924–1937 (2007).

24. L. Lombardelli et al., HLA-G5 induces IL-4 secretion critical for successful pregnancy through differential expression of ILT2 receptor on decidual CD4(+) T cells and macrophages. J Immunol 191, 3651–3662 (2013).

25. S. Endo, Y. Sakamoto, E. Kobayashi, A. Nakamura, T. Takai, Regulation of cytotoxic T lymphocyte triggering by PIR-B on dendritic cells. Proc Natl Acad Sci U S A 105, 14515–14520 (2008).

26. S. Pereira, H. Zhang, T. Takai, C. A. Lowell, The inhibitory receptor PIR-B negatively regulates neutrophil and macrophage integrin signaling. J Immunol 173, 5757–5765 (2004).

27. N. S. Wilson et al., An Fcgamma receptor-dependent mechanism drives antibody-mediated target-receptor signaling in cancer cells. Cancer Cell 19, 101–113 (2011).

28. E. Soderlind et al., Recombining germline-derived CDR sequences for creating diverse single-framework antibody libraries. Nat Biotechnol 18, 852–856 (2000).

29. A. Roghanian et al., Antagonistic human FcgammaRIIB (CD32B) antibodies have anti-tumor activity and overcome resistance to antibody therapy in vivo. Cancer Cell 27, 473–488 (2015).

30. D. C. Jones et al., Allele-specific recognition by LILRB3 and LILRA6 of a cytokeratin 8-associated ligand on necrotic glandular epithelial cells. Oncotarget 7, 15618–15631 (2016).

31. D. Saverino et al., The CD85/LIR-1/ILT2 inhibitory receptor is expressed by all human T lymphocytes and down-regulates their functions. J Immunol 165, 3742–3755 (2000).

32. M. Shiroishi et al., Human inhibitory receptors Ig-like transcript 2 (ILT2) and ILT4 compete with CD8 for MHC class I binding and bind preferentially to HLA-G. Proc Natl Acad Sci U S A 100, 8856–8861 (2003).

33. C. C. Chang et al., BCL6 Is Required for Differentiation of Ig-Like Transcript 3-Fc-Induced CD8(+) T Suppressor Cells. J Immunol 185, 5714–5722 (2010).

34. K. Hussain et al., Upregulation of FcgammaRIIb on monocytes is necessary to promote the superagonist activity of TGN1412. Blood 125, 102–110 (2015).

35. L. N. Dahal, A. Roghanian, S. A. Beers, M. S. Cragg, FcgammaR requirements leading to successful immunotherapy. Immunol Rev 268, 104–122 (2015).

36. A. Roghanian et al., Cyclophosphamide Enhances Cancer Antibody Immunotherapy in the Resistant Bone Marrow Niche by Modulating Macrophage FcgammaR Expression. Cancer Immunol Res 7, 1876–1890 (2019).

37. I. Leskov et al., Rapid generation of human B-cell lymphomas via combined expression of Myc and Bcl2 and their use as a preclinical model for biological therapies. Oncogene 32, 1066–1072 (2013).

38. F. Y. McWhorter, T. Wang, P. Nguyen, T. Chung, W. F. Liu, Modulation of macrophage phenotype by cell shape. Proc Natl Acad Sci U S A 110, 17253–17258 (2013).

39. S. Galvan-Pena, L. A. O’Neill, Metabolic reprograming in macrophage polarization. Front Immunol 5, 420 (2014).

40. R. C. Khanolkar et al., Leukocyte Ig-Like receptor B1 restrains dendritic cell function through increased expression of the NF-kappaB regulator ABIN1/TNIP1. J Leukoc Biol, (2016).

41. Y. Zhao et al., The Orphan Immune Receptor LILRB3 Modulates Fc Receptor-Mediated Functions of Neutrophils. J Immunol 204, 954–966 (2020).

42. D. E. Sloane et al., Leukocyte immunoglobulin-like receptors: novel innate receptors for human basophil activation and inhibition. Blood 104, 2832–2839 (2004).

43. A. A. Bashirova et al., Diversity of the human LILRB3/A6 locus encoding a myeloid inhibitory and activating receptor pair. Immunogenetics 66, 1–8 (2014).

44. M. D. Pescovitz, Rituximab, an anti-cd20 monoclonal antibody: history and mechanism of action. Am J Transplant 6, 859–866 (2006).

45. X. Yu et al., Complex Interplay between Epitope Specificity and Isotype Dictates the Biological Activity of Anti-human CD40 Antibodies. Cancer Cell 33, 664–675 e664 (2018).

46. S. Liang, W. Zhang, A. Horuzsko, Human ILT2 receptor associates with murine MHC class I molecules in vivo and impairs T cell function. Eur J Immunol 36, 2457–2471 (2006).

47. W. Zhang, S. Liang, J. Wu, A. Horuzsko, Human Inhibitory Receptor ILT2 Amplifies CD11b(+)Gr1(+) Myeloid-Derived Suppressor Cells that Promote Long-Term Survival of Allografts. Transplantation 86, 1125–1134 (2008).

48. M. Nakayama et al., Paired Ig-like receptors bind to bacteria and shape TLR-mediated cytokine production. J Immunol 178, 4250–4259 (2007).

49. J. Zheng et al., Inhibitory receptors bind ANGPTLs and support blood stem cells and leukaemia development. Nature 485, 656–660 (2012).

50. F. O. Martinez, S. Gordon, M. Locati, A. Mantovani, Transcriptional profiling of the human monocyte-to-macrophage differentiation and polarization: new molecules and patterns of gene expression. J Immunol 177, 7303–7311 (2006).

51. D. M. W. Zaiss, W. C. Gause, L. C. Osborne, D. Artis, Emerging functions of amphiregulin in orchestrating immunity, inflammation, and tissue repair. Immunity 42, 216–226 (2015).

52. Y. L. Hsu et al., Lung tumor-associated dendritic cell-derived amphiregulin increased cancer progression. J Immunol 187, 1733–1744 (2011).

53. P. Vlaicu et al., Monocytes/macrophages support mammary tumor invasivity by co-secreting lineage-specific EGFR ligands and a STAT3 activator. BMC Cancer 13, 197 (2013).

54. B. Busser, L. Sancey, E. Brambilla, J. L. Coll, A. Hurbin, The multiple roles of amphiregulin in human cancer. Biochim Biophys Acta 1816, 119–131 (2011).

55. W. van der Touw et al., Glatiramer Acetate Enhances Myeloid-Derived Suppressor Cell Function via Recognition of Paired Ig-like Receptor B. J Immunol 201, 1727–1734 (2018).

56. H. M. Chen et al., Blocking immunoinhibitory receptor LILRB2 reprograms tumor-associated myeloid cells and promotes antitumor immunity. J Clin Invest, (2018).

57. M. Deng et al., LILRB4 signalling in leukaemia cells mediates T cell suppression and tumour infiltration. Nature 562, 605–609 (2018).

58. N. Olsson et al., Proteomic Analysis and Discovery Using Affinity Proteomics and Mass Spectrometry. Mol Cell Proteomics 10, (2011).

59. A. Roghanian et al., Filament-associated TSGA10 protein is expressed in professional antigen presenting cells and interacts with vimentin. Cellular immunology 265, 120–126 (2010).

60. L. E. Hogan, D. C. Jones, R. L. Allen, Expression of the innate immune receptor LILRB5 on monocytes is associated with mycobacteria exposure. Sci Rep 6, 21780 (2016).

61. B. Langmead, C. Trapnell, M. Pop, S. L. Salzberg, Ultrafast and memory-efficient alignment of short DNA sequences to the human genome. Genome Biol 10, R25 (2009).

62. B. Li, C. N. Dewey, RSEM: accurate transcript quantification from RNA-Seq data with or without a reference genome. BMC Bioinformatics 12, 323 (2011).

63. M. D. Robinson, D. J. McCarthy, G. K. Smyth, edgeR: a Bioconductor package for differential expression analysis of digital gene expression data. Bioinformatics 26, 139–140 (2010).

64. D. W. Huang et al., DAVID Bioinformatics Resources: expanded annotation database and novel algorithms to better extract biology from large gene lists. Nucleic Acids Res 35, W169–175 (2007).

65. A. Subramanian et al., Gene set enrichment analysis: a knowledge-based approach for interpreting genome-wide expression profiles. Proc Natl Acad Sci U S A 102, 15545–15550 (2005).

66. A. I. Saeed et al., TM4: a free, open-source system for microarray data management and analysis. Biotechniques 34, 374–378 (2003).

